# Novel leaf-root coordination driven by leaf water storage tissues in mangroves

**DOI:** 10.1101/2022.07.26.501578

**Authors:** Jingjing Cao, Qingpei Yang, Jing Chen, Mingzhen Lu, Weizheng Ren, Yanmei Xiong, Yuxin Pei, Deliang Kong

**Author notes:** **Correspondence and requests for materials should be addressed to D.K. (**).

## Abstract

Interactions among root and leaf traits (in particular, leaf hydraulic and leaf economics traits) are fundamental in generating diverse strategies in land plants, yet it remains a knowledge gap in mangrove plants that experiences saline stress distinct from most other vascular plants. Here, we tested the trait relationships in mangrove plants and compared them with typical land plants (non-mangrove). Consistent with non-mangrove plants, leaf hydraulic and economics traits were decoupled in mangrove plants. However, mangrove leaf economics traits correlated strongly with root hydraulic traits, which are normally decoupled in non-mangrove plants. Moreover, we observed a unique scaling relationship between *leaf dry mass per area* and root hydraulic traits in mangroves. The novel coordination between leaves and roots arises from the wide-presence of leaf *water storage tissues* in mangroves, and this potentially represents a new paradigm with which we look into the ecology, physiology and evolution of this important vegetation.

## Introduction

Photosynthesis, the conversion of atmospheric CO_2_ into carbohydrates by the leaves using light, is the most important biological process on Earth. During this process, atmospheric CO_2_ enters the leaf mesophyll, mainly the palisade cells, and is fixed as carbohydrates through a series of biochemical reactions^1^. Researchers coined a term, “leaf economics spectrum”, to describe a tradeoff in this process across species worldwide; that is, leaves with higher leaf CO_2_ fixation rates have a shorter leaf lifespan, and vice versa^2,3,4,5^.

The maintenance of photosynthesis relies greatly on water supply by leaf veins, and a large portion of water is lost by transpiration through stomata. The well-correlated vein and stomatal traits depicting water supply into and loss out of leaves constitute leaf hydraulic traits, or the leaf hydraulics. While leaf hydraulics is vital to the process of leaf photosynthesis (which lead us to hypothesize a coupling of leaf hydraulics and economics), recent findings however suggest that leaf hydraulics is decoupled from leaf economics across a range of non-mangrove plants^6,7^. This counterintuitive finding thus further suggests that leaves should be understood by looking at multiple trait dimensions (often referred to as “multidimensionality”). The multidimensionality in leaves uncovers a ubiquitous mechanism by which species coexist and respond to global climate change^7^.

The decoupling between leaf hydraulics and economics can essentially arise from how the water is partitioned between leaf photosynthesis and transpiration, although previously studies ascribed such decoupling to functional modularity of leaf anatomic structures (i.e., mesophyll and veins), non-simultaneous evolution of leaf mesophyll and leaf veins, and selection by environmental heterogeneity^7^. Specifically, plants generally lose hundreds, even thousands, of mol of water through transpiration in exchange for the fixation of 1 mol of atmospheric CO_2_^1^. Theoretically, 1 mol water is consumed in the biochemical fixation of 1 mol CO_2,_ and we refer to such water as “*water used for photosynthetic metabolism* (*WUPM*).” Undoubtedly, *WUPM* is closely related to leaf economics, whereas “*water used for transpiration* (termed as *WUT*) ” is tightly associated with leaf hydraulic traits^7,8^. Given that *WUPM* is negligible compared to *WUT*, it is likely that *WUPM* and hence leaf economics could have little impact on leaf hydraulics. Therefore, the contrasting amount and functioning of *WUPM* and *WUT* constitute a novel mechanism that essentially drives leaf economics-hydraulics decoupling^5,7,9^.

However, our current knowledge on the relationship between leaf hydraulics and economics comes mainly from land plants. It remains unclear whether the decoupling between leaf hydraulics and economics still holds and whether the above novel mechanism accounting for such decoupling can be applied to plants under stressful environments such as coastal mangrove species that typically grow under high salinity and are confronted with severe physiological drought^5,10,11^. One conspicuous characteristic of mangrove plants is that they usually have significant *water storage tissues* in their leaves as an adaption to the imperative water demanding of transpiration under drought stress^12,13,14,15,16^. Meanwhile, plants usually adjust above- and belowground organs (e.g., leaves vs. roots) in a coordinated manner during plant evolution and response to changing environments^4,17,18,19,20,21^. However, little is known about how leaves, especially leaf *water storage tissues*, and roots vary across mangrove species and how above- and belowground coordination, if present, differs from that of non-mangrove plants.

The absorption and transport of water under strong osmotic stress due to high soil salinity are crucial for growth and survival of mangrove plants^22,23,24,25^. Root anatomical structures, such as the stele and cortex, are directly related to root water absorption and transportation^26,27,28^, and the anatomical traits, hereafter termed root hydraulic traits or root hydraulics. For root hydraulics in non-mangrove plants, the *thickness of root tissues outside the stele* (including epidermis, exodermis and cortex) has been widely observed to increase considerably faster than the stele radius with increasing root diameter^26,29,30^. This allometry in root structures is insightful for understanding plant evolution and responses to drought and carbon (C) limitation under stress^31,32,33^ that mangrove species are exposed to in their natural habitats. However, it is still unclear whether allometry between *root tissues outside the stele* and stele exists in mangrove species and how such allometry, if present, cooperates with leaves in adaptation to salinity stress.

In the present study, we selected 17 representative mangrove species in a reserve in southern China in which the most diverse mangrove species in China are found. We measured 10 leaf hydraulic and economics traits, including leaf *water storage tissues*, and 4 root hydraulic traits. To compare trait relationships between mangrove and non-mangrove plants, we constructed a dataset of land species with leaf and root traits measured concomitantly ^7,26^. We tested the following hypotheses: (1) compared with the independence between leaf hydraulics and economics in non-mangrove plants, the two trait spectrum should be coupled in mangrove plants under high-salinity selection pressure; (2) in mangrove plants, root hydraulics should be positively correlated with both leaf hydraulics and economics, which would be different from non-mangrove plants where root hydraulics is positively correlated with leaf hydraulics^18,28^, yet decoupled from leaf economics.

## Results

### Traits association between leaf economics and leaf hydraulics

The first principal component analysis (PCA) axis explained 52.53% of total leaf trait variation in leaves of mangrove species. Leaf economics traits, such as *leaf dry mass per area, specific leaf area, leaf mass-based nitrogen concentration, leaf thickness*, leaf *water storage tissues*, and *total phenol content*, had relatively high loading scores on this axis (Fig. 1a, Table S1). *Leaf*

*mass-based nitrogen concentration* was negatively correlated with *leaf dry mass per area* (*r* = -0.86, *p* < 0.001) and leaf *water storage tissues* was positively correlated with *leaf dry mass per area* and *leaf thickness* (*r* = 0.71, *p* = 0.001; *r* = 0.62, *p* = 0.008) (Table S2). The second PCA axis accounted for 21.28% of the total leaf trait variation. *Leaf minor vein density, leaf minor vein diameter, leaf tissue density*, and *leaf C isotope composition* (mainly hydraulic traits) had high loading on this axis (Fig. 1a, Table S1). *Leaf minor vein diameter* was negatively correlated with *leaf minor vein density* (*r* = -0.90, *p* < 0.001); neither vein trait was correlated with *leaf tissue density*, or *leaf C isotope composition* (Table S2). In addition, permutation analysis of mangrove plants showed a significant coupling of leaf *water storage tissues* with leaf economics (*p* = 0.003) instead of leaf hydraulics (*p* = 0.338) (Fig. S1).

**Figure 1.**
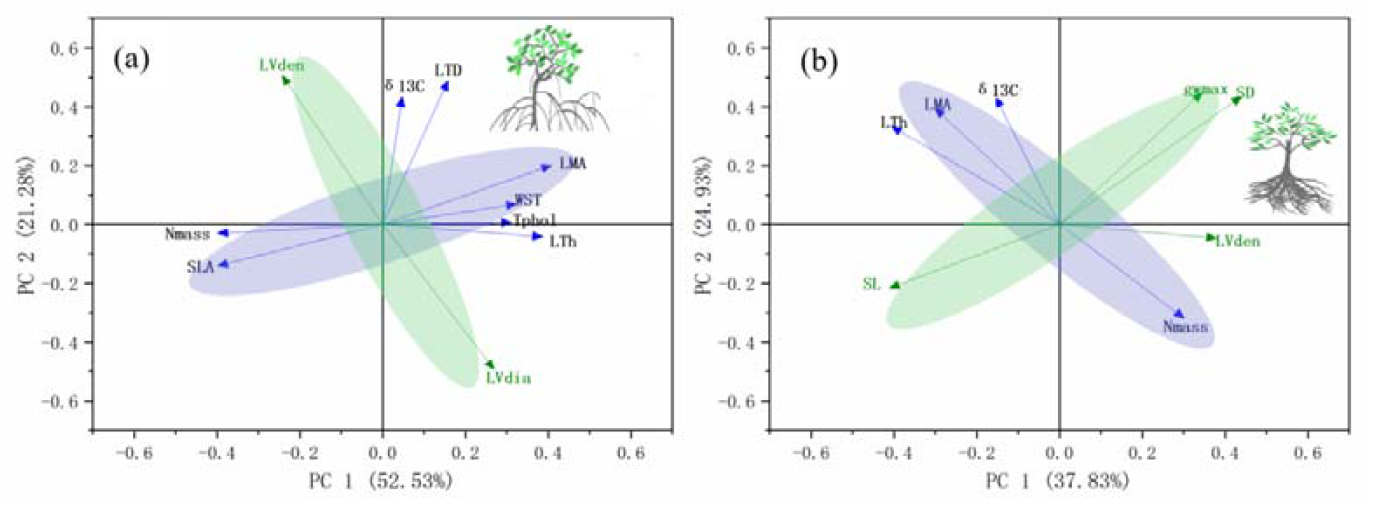
Traits decoupling of leaf economics and hydraulics shared among mangrove and non-mangrove plants. (**a**) In mangrove trees, leaf economics traits (denoted in blue; e.g., LMA, SLA) do not align with leaf hydraulics traits (denoted in green; e.g., LV_dia_) according to a principal component analysis of 17 species, suggesting the decoupling of the leaf economics and hydraulics. (**b**) In non-mangrove plants, we observe a similar pattern of traits decoupling between leaf economics and leaf hydraulics, suggested by the largely orthogonal alignment between the two groups of traits (blue vs. green). Traits abbreviation: LTh, *leaf thickness*; LMA, *leaf dry mass per area*; N_mass_, *leaf mass-based nitrogen concentration*; δ^13^C, *leaf carbon isotope composition*; SLA, *specific leaf area*; LTD, *leaf tissue density*; Tphol, *total phenol content*; WST, *water storage tissues*; LV_dia_, *leaf minor vein diameter*; LV_den_, *leaf minor vein density*; SL, *stomatal guard cell length*; SD, *stomatal density*; g_wmax,_ *maximum stomatal conductance to water vapor*.

Regarding non-mangrove species, the first PCA axis of leaves explained 37.83% of total leaf trait variation. *Leaf thickness, leaf minor vein density, stomatal guard cell length, stomatal density*, and *maximum stomatal conductance to water vapor* had relatively high loading scores on this axis (Fig. 1b, Table S3). Among leaf hydraulic traits, *leaf minor vein density* was negatively correlated with *stomatal guard cell length* (*r* = -0.48, *p* < 0.001) and positively correlated with *stomatal density* (*r* = 0.36, *p* = 0.001) (Table S4). The second PCA axis accounted for 24.93% of the total leaf trait variation. *Leaf dry mass per area*, and *leaf mass-based nitrogen concentration, leaf C isotope composition, stomatal density*, and *maximum stomatal conductance to water vapor* had high loading scores on this axis (Fig. 1b, Table S3). Among leaf economics traits, *leaf dry mass per area* was negatively correlated with *leaf mass-based nitrogen concentration* (*r* = -0.49, *p* < 0.001) (Table S4).

### Leaf-root traits coordination

When leaf and root traits were pooled in mangrove plants, the first two axes of the leaf-root combined PCA accounted for 50.73% and 15.87% of the total trait variation, respectively (Fig. 2a). Root and leaf traits with high loading scores separately in the first axis of the root and the leaf PCA also had high loading scores in the first axis of the leaf-root combined PCA. *Leaf minor vein density* and *minor vein diameter* had high loading scores in the second axis of the combined PCA (Fig. 2a, Table S5).

**Figure 2.**
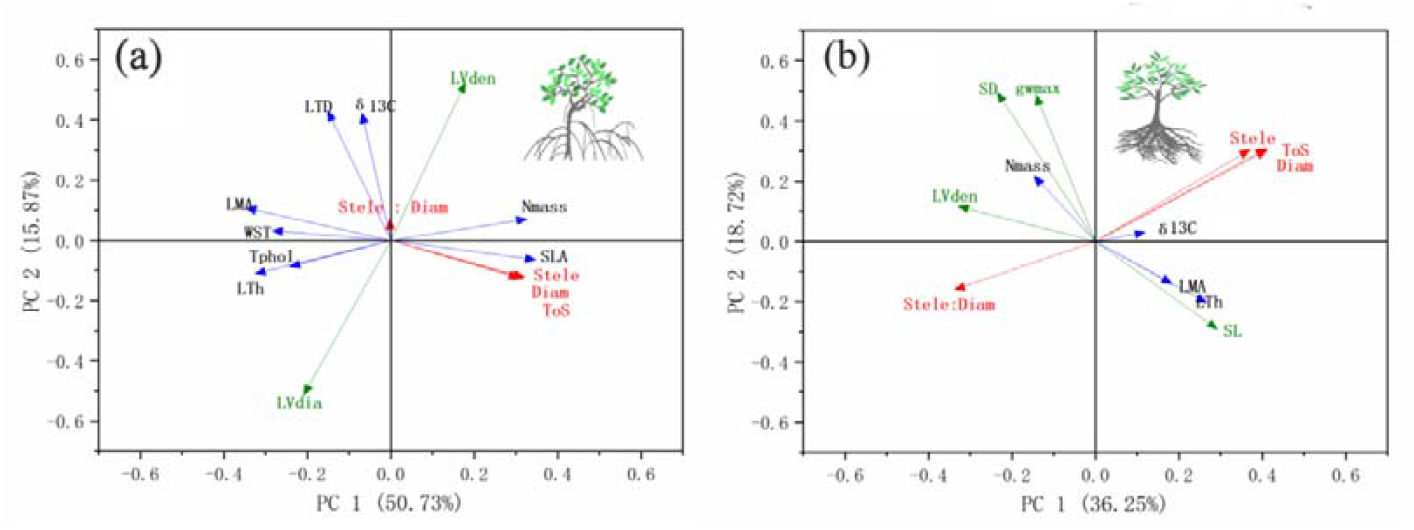
Leaf-root traits coordination differs between mangrove and non-mangrove plants. (**a**) In mangrove trees, the newly added root traits (in red), align with majority of leaf economics traits (denoted in blue consistent with Fig.1; e.g., LMA) and largely orthogonal to leaf hydraulic traits (denoted in green consistent with Fig.1). (**b**) In contrast with mangrove trees, root traits of non-mangrove plants appear to be orthogonal to leaf economics traits. Abbreviations of the traits are: LTh, *leaf thickness*; LMA, *leaf dry mass per area*; N_mass_, *leaf mass-based nitrogen concentration*; δ^13^C, *leaf carbonisotope composition*; SLA, *specific leaf area*; LTD, *leaf tissue density*; Tphol, *total phenol content*; WST, *water storage tissues*; LV_dia_, *leaf minor vein diameter*; LV_den_, *leaf minor vein density*; Diam, *root diameter*; ToS, *thickness of root tissues outside the stele*; Stele, *root stele diameter*; Stele : Diam, *stele to root diameter ratio*; SL, *stomatal guard cell length*; SD, *stomatal density*; g_wmax,_ *maximum stomatal conductance to water vapor*.

In non-mangrove species, the first two axes of the leaf-root combined PCA accounted for 36.25% and 18.72% of the total trait variation, respectively. *Leaf minor vein density, root diameter, thickness of root tissues outside the stele*, and *stele to root diameter ratio* had high loading scores in the first axis of the combined PCA. S*tomatal density, maximum stomatal conductance to water vapor, thickness of root tissues outside the stele*, and *root stele diameter* had high loading scores in the second axis of the combined PCA (Fig. 2b, Table S6).

Permutation analysis of the first PCA scores of the three trait groups (i.e., leaf hydraulic traits, leaf economics traits, and root hydraulic traits, Fig. 2a) showed independence of leaf hydraulics from both leaf economics and root hydraulics (*p* = 0.117, *p* = 0.218, respectively) In contrast leaf economics was coupled with root hydraulics in mangrove species (*p* < 0.001) (Fig. 2a, Fig. S2, Table S5). In the 78 non-mangrove plant species, leaf hydraulics, although decoupled from leaf economics (*p* = 0.062), correlated with root hydraulics (*p* = 0.012), and the leaf economics of these species were orthogonal to root hydraulics (*p* = 0.073) (Fig. 2b, Fig. S3, Table S6). Overall, the effect of phylogeny on leaf and root traits was relatively less pronounced in mangrove plants than in non-mangrove plants (Tables 1 and 2), suggesting the importance of environmental selection in shaping mangrove plant traits. Phylogenetically independent contrast-based relationships among leaf hydraulics, economics, and root hydraulics were generally similar to the results obtained from mangrove plants’ original trait data (Fig. S4, S5).

**Table 1.** Variation in 14 root and leaf functional traits measured from17 mangrove species.

|  | Functional traits | Unit | Mean | SE* | CV† | Max | Min | Blomberg's K |
| --- | --- | --- | --- | --- | --- | --- | --- | --- |
| Leaf | *Leaf thickness* | mm | 0.42 | 0.036 | 34.93 | 0.65 | 0.13 | 0.17 |
|  | *Leaf dry mass per area* | g cm<sup>-2</sup> | 0.01 | 0.001 | 39.31 | 0.02 | 0.00 | 0.12 |
|  | *Leaf mass-based nitrogen concentration* | mg g<sup>-1</sup> | 2.01 | 0.181 | 37.13 | 3.95 | 1.09 | 0.10 |
|  | *Leaf carbon isotope composition* | ‰ | -30.58 | 0.397 | -5.35 | -27.01 | -32.93 | 0.08 |
|  | *Specific leaf area* | cm<sup>2</sup> g<sup>-1</sup> | 92.94 | 13.424 | 59.55 | 245.60 | 40.41 | 0.17 |
|  | *Leaf tissue density* | g cm<sup>-3</sup> | 0.32 | 0.013 | 17.07 | 0.39 | 0.20 | 0.08 |
|  | *Total phenol content* | % | 2.34 | 0.258 | 45.42 | 4.37 | 0.60 | 0.08 |
|  | *Water storage tissues* | % | 0.13 | 0.014 | 43.39 | 0.24 | 0.06 | 0.08 |
|  | *Leaf minor vein diameter* | µm | 49.13 | 8.922 | 74.88 | 156.64 | 15.50 | 0.08 |
|  | *Leaf minor vein density* | mm mm<sup>-2</sup> | 10.66 | 2.383 | 92.13 | 42.85 | 2.30 | 0.10 |
| Root | *Root diameter* | µm | 191.01 | 18.300 | 39.50 | 388.03 | 112.24 | 0.26 |
|  | *Thickness of root tissues outside the stele* | µm | 78.19 | 7.826 | 41.26 | 163.32 | 44.11 | **0.26** |
|  | *Root stele diameter* | µm | 34.28 | 3.232 | 38.88 | 61.42 | 20.19 | 0.17 |
|  | *Stele to root diameter ratio* | % | 0.19 | 0.009 | 20.20 | 0.24 | 0.12 | 0.08 |

Root traits were measured for the first-order roots.
SE\*, standard error.
CV†, coefficient of variance.
Blomberg's K values in bold indicate they are significant at $p < 0.05$ .

**Table 2.** Variation in 12 root and leaf functional traits measured from 78 non-mangrove plant species.

|  | Functional traits | Unit | Mean | SE* | CV† | Max | Min | Blomberg's K |
| --- | --- | --- | --- | --- | --- | --- | --- | --- |
| Leaf | *Leaf thickness* | mm | 0.24 | 0.01 | 32.65 | 0.56 | 0.09 | 0.01 |
|  | *Leaf dry mass per area* | g m<sup>-2</sup> | 79.53 | 2.89 | 32.72 | 159.17 | 32.53 | 0.01 |
|  | *Leaf mass-based nitrogen concentration* | mg g<sup>-1</sup> | 17.70 | 0.69 | 35.30 | 36.28 | 5.40 | 0.01 |
|  | *Leaf carbon isotope composition* | ‰ | -31.93 | 0.21 | -5.93 | -27.38 | -36.72 | **0.22** |
|  | *Leaf minor vein density* | mm mm<sup>-2</sup> | 6.12 | 0.22 | 32.22 | 13.01 | 2.28 | 0.01 |
|  | *Stomatal guard cell length* | µm | 20.97 | 0.57 | 24.61 | 35.40 | 9.56 | **0.35** |
|  | *Stomatal density* | mm<sup>-2</sup> | 398.79 | 29.85 | 67.37 | 1716.07 | 91.47 | 0.04 |
|  | *Maximum stomatal conductance to water vapor* | mol mm<sup>-2</sup> s<sup>-1</sup> | 1.45 | 0.08 | 47.40 | 4.83 | 0.37 | 0.02 |
| Root | *Root diameter* | µm | 362.98 | 22.82 | 56.58 | 1010.00 | 73.00 | **0.42** |
|  | *Thickness of root tissues outside the stele* | µm | 232.36 | 17.27 | 66.89 | 675.00 | 26.00 | **0.22** |
|  | *Root stele diameter* | µm | 83.28 | 4.74 | 51.25 | 258.00 | 24.00 | 0.07 |
|  | *Stele to root diameter ratio* | % | 0.25 | 0.01 | 20.96 | 0.39 | 0.16 | 0.02 |

Root traits were measured for the first-order roots.
SE\*, standard error.
CV†, coefficient of variance.
Blomberg's K values in bold indicate they are significant at $p < 0.05$ .

### Leaf-root allometry

The root stele radius and *thickness of root tissues outside the stele* followed an allometric relationship with increasing *root diameter* in both mangrove and non-mangrove species (Fig. S6). As the keystone trait of leaves, *leaf dry mass per area* (LMA) is perhaps the single most important organizing trait among all other leaf traits. Surprisingly, we found a strong allometric relationship between LMA and root anatomical traits only in mangrove plants (*p* < 0.001, each; Fig. 3a). Specifically, the regression slope (in absolute values) of the relationship between *thickness of root tissues outside the stele* and LMA was higher than the regression slope of the relationship between stele radius and LMA (standardized major axis, *p* < 0.001). Leaf-root allometry was still detected when other leaf economics traits, such as *leaf thickness, specific leaf area*, leaf *water storage tissues*, and *leaf mass-based nitrogen concentration* were used (Fig. S7a–d). Unlike mangrove plants, neither of the above two root anatomical traits were correlated with LMA in non-mangrove species (Fig. 3b).

**Figure 3.**
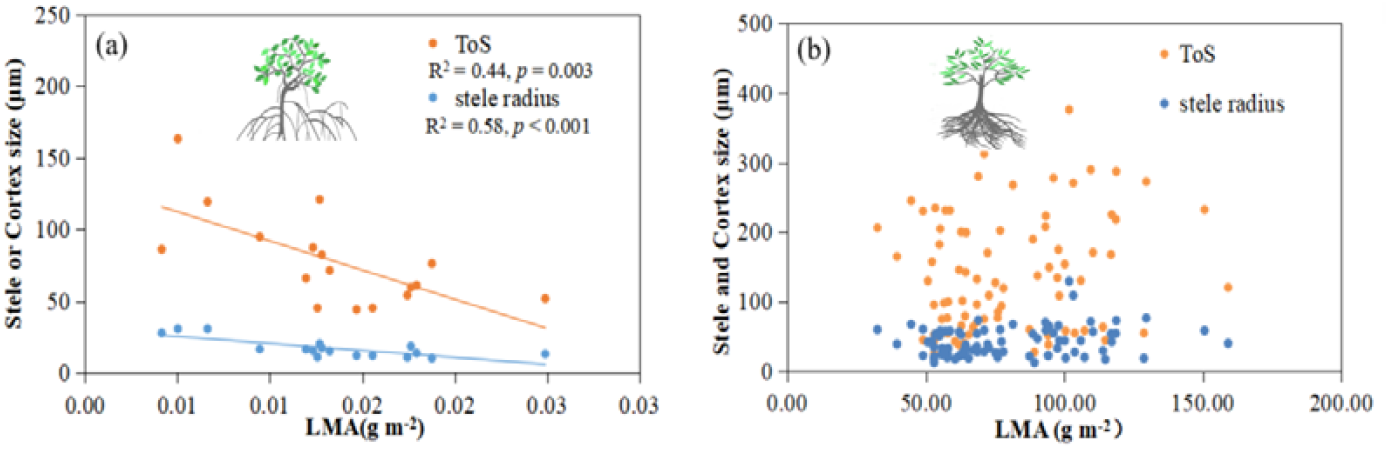
Allometric relationships between roots and leaves in mangrove (a) and non-mangrove plants (b). Root anatomic structures refer to root stele radius and *thickness of root tissues outside the stele*. Abbreviations: ToS, *thickness of root tissues outside the stele*; Diam, *root diameter*; LMA, *leaf dry mass per area*.

## Discussion

### Decoupling of leaf economics from leaf hydraulics shared among mangrove and non-mangrove plants

Contrary to our first hypothesis postulating coupled leaf economics with leaf hydraulics in mangrove plants, the two trait spectra were decoupled in mangrove leaves (Fig. 1a, Fig. S2a) as has been commonly observed in non-mangrove plants. Unlike most non-mangrove plants, mangrove plants usually have remarkable *water storage tissues* in their leaves to adapt to physiological drought caused by high-salinity environments^34^. For example, it has been reported that the mangrove plant *Avicennia marina* can maintain a normal transpiration rate for up to 2 h under water supply from leaf *water storage tissues* without water supply from roots^35,36^. Therefore, leaf *water storage tissues* is important for water that is required for leaf transpiration in mangrove plants^35,37,38^, which may, as articulated below, underlie the decoupling of leaf economics and hydraulics in mangrove leaves.

Under high soil salinity, plant leaves usually suffer greatly from water deficiency in transpiration due to insufficient water supply from the roots^5,11^. For example, in the mangrove species *Ceriops tagal*, the vessel diameter of absorptive roots under high salinity is only two-thirds of that under low salinity. Theoretically, this leads to root water-supply efficiency under high salinity only 20% as that under low salinity^5^. In coping with such water deficiency, mangrove plants can increase the amount of water stored in leaf *water storage tissues* for one hand^35,36,39^, and for the other hand speed up the release of water from the *water storage tissues* for transpiration^37^. Assuming no such water supply from leaf *water storage tissues*, photosynthetic cells in the palisade would be dehydrated, and photosynthesis would then be greatly inhibited^36^. Therefore, leaf *water storage tissues* could be closely related to leaf photosynthesis and leaf economics, given their functioning as an agent to protect photosynthetic cells from damage caused by dehydration. The physiologically based coupling of leaf *water storage tissues* with leaf economics also well explains their statistical coupling (Fig. S1).

Here, we used mangroves under high salinity as an example to understand the role of leaf *water storage tissues* in shaping the decoupling relationship between leaf hydraulics and economics. Under such salinity stress, mangrove leaf hydraulic traits, such as, *leaf minor vein diameter*, are faced with two competing selection pressures: (1) reducing *leaf minor vein diameter* due to inadequate water supply from thinner root vessels^5,11,40^; and (2) increasing the size of leaf *water storage tissues*^35^ entails a thick *leaf minor vein diameter* to meet the water requirement of the increased leaf *water storage tissues* (Fig. 4). Therefore, the variation in *leaf minor vein diameter* with increasing soil salinity may be markedly attenuated by the two opposing section pressures, whereas the size of leaf *water storage tissues* tended to increase in high salinity. The different trajectories of leaf hydraulics (exemplified by *leaf minor vein diameter*) and leaf *water storage tissues* with soil salinity combined with the coupling between *water storage tissues* and leaf economics may explain the independence between leaf hydraulics and economics.

**Figure 4.**
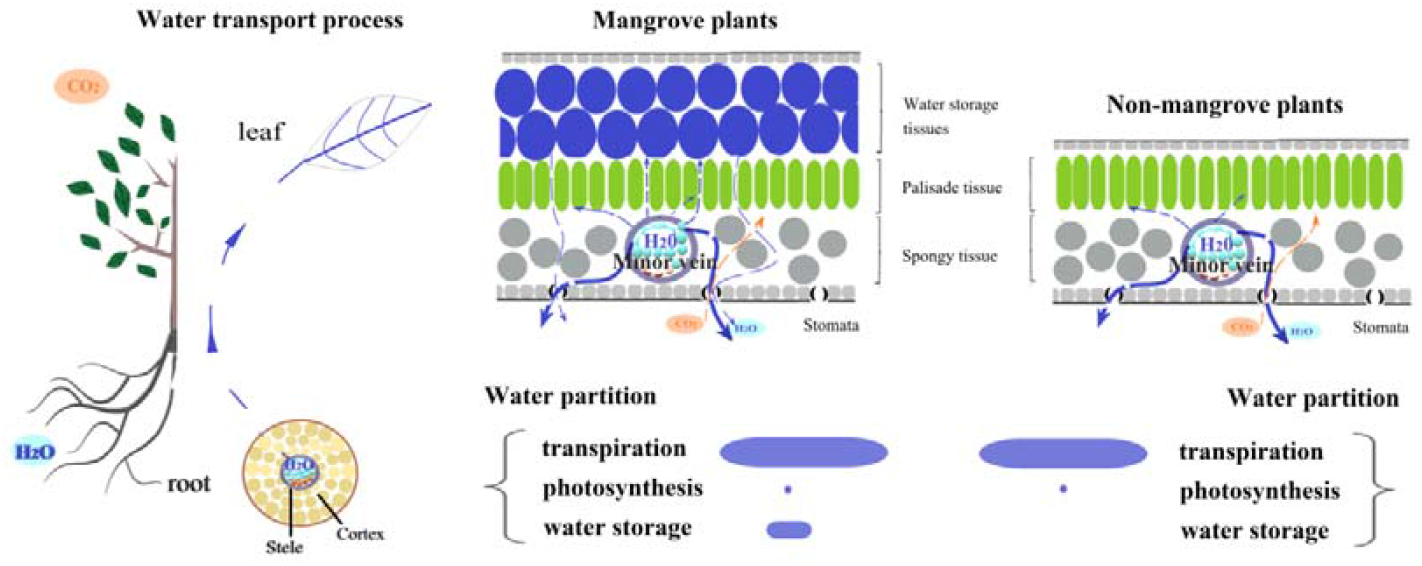
A conceptual framework of water transport and partition from roots to leaves for mangrove and non-mangrove plants. The blue dashed lines in leaves represent three ways of water partition from minor veins to (1) water storage tissue in leaves and loss from stomata, (2) palisade tissue for photosynthesis, and (3) transpiration via stomata in mangrove plants; water partition ways in non-mangrove plants usually include the above (2) and (3). The CO_2_ fixation and hence the photosynthate are indicated with solid circles and dashed lines in orange. The relative amount of water used in the above partition ways, by referring to studies on mangroveplants^35,36^, was roughly represented by different-sized dashed lines and stripes in blue.

In mangrove plants, leaf *water storage tissues* usually stores a considerable amount of water^35,36^. After loss through transpiration, leaf *water storage tissues* can be rehydrated rapidly through various sources, such as rainfall, dew, fog, and root water supply^38^. Frequent water loss–rehydration cycling suggests that leaf *water storage tissues* in mangroves contribute considerably contribution to transpiration. Therefore, the quantity of water associated with leaf economics (i.e., *WUPM* + water storage in leaf *water storage tissues*) may be comparable, rather than negligible, to the amount of water used for transpiration (Fig. 4), and leaf economics and hydraulics in mangrove plants should be coupled according to the novel mechanism proposed in the Introduction section. Contrary to this expectation, leaf economics and hydraulics were decoupled in mangroves. The decoupling of leaf economics and hydraulics in mangroves could arise primarily from the effect of the selection pressure of leaf *water storage tissues*, as a key component of leaf economics, on the leaf hydraulic traits (e.g., *leaf minor vein diameter*). This also suggests that our proposed novel mechanism that drives decoupled leaf economics and hydraulics may hold only for land plants that do not possess leaf *water storage tissues*, as such lacking selection pressure on leaf hydraulic traits.

### Leaf-root traits coordination differs between mangrove and non-mangrove plants

In mangrove plants, root hydraulics had a significant correlation with leaf economics but no correlation with leaf hydraulics (Fig. 2a, Fig. S2). This is partially consistent with our second hypothesis for mangrove plants. Generally, under drought or physiological drought caused by high salinity, plant roots and vessels tend to be thinner^11,21,41,42,43^, whereas leaf *water storage tissues* usually increases ^34,35,39^. These contrasting trends will cause a negative relationship between root diameter (representative of root hydraulics) and leaf *water storage tissues* (representative of leaf economics) in mangrove plants, which was confirmed by our results (Fig. 2a, Table S5). Given that leaf hydraulics, as argued previously, is decoupled from leaf *water storage tissues*, it is reasonable to observe a decoupling relationship between root hydraulics and leaf hydraulics in mangroves (Fig. 2a, Fig. S2b).

In contrast to mangrove plants, root hydraulics in non-mangrove plants is well-coordinated with leaf hydraulics and decoupled from leaf economics (Fig. 2b, Fig. S3). The significant relationship between root and leaf hydraulics may be because they belong to a common and continuous vessel system across above- and belowground plant parts. Owing to the decoupled relationship between leaf hydraulics and economics^6,7^, it can be deduced that root hydraulics is decoupled from leaf economics in non-mangrove plants. Such expected root-leaf relationships have also been confirmed in recent studies, for example, significant relationships between root hydraulics (represented by *root diameter*) and *leaf minor vein density*^18^ and leaf transpiration^28^, but no relationship with leaf economics^19^. Obviously, non-mangrove and mangrove plants diverge greatly with regard to in the root-leaf relationships. Such divergence may be attributed to the widespread presence of leaf *water storage tissues* in mangrove plants, which is absent in most non-mangrove plants.

### Above-belowground trait coordination explained by leaf-root allometric relationship

Allometry between the *root tissues outside the stele* and the stele in mangrove plant roots, similar to that in non-mangrove plant roots, was revealed for the first time in this study (Fig. S6a, b). Allometry in mangrove roots, as is the case in non-mangrove plants, could be formed to achieve a balance between nutrient absorption and transport as well as a balance between C supply and consumption^32,44^. In addition to allometry within roots, we found a conspicuous allometry between roots and leaves in mangrove plants. The *thickness of root tissues outside the stele* increased faster than the stele radius with decreasing *leaf dry mass per area* (Fig. 3a). In contrast, no such allometry was observed in non-mangrove plants (Fig. 3b).

The contrasting leaf-root allometry between mangrove and non-mangrove plants could be related to the difference in C cycling within plants. Specifically, the well-recognized decoupling between leaf economics and hydraulics can also convey an independence between C production and transportation because leaf veins function as C transportation via sieves besides water transportation via vessels, and the sieves and vessels are closely related in structure and functioning in land plants^32,45^. Physiologically, such decoupling between C production and transportation can be explained by different environmental factors driving these processes. For example, leaf C production depends greatly on light, whereas C transportation is governed by water and phosphorus availability^1^; these environmental factors always vary independently^5,7,46^.

Similar to leaf veins, the *root tissues outside the stele* and the stele in absorptive roots, although previously treated as hydraulic traits, also serve as key agents for C supply (through sieves in the stele) and C consumption (through *root tissues outside the stele* cells), respectively. The allometry between the *root tissues outside the stele* and the stele is thought to arise from a balance between C supply and consumption within the roots^32^. Since C supply in roots and C transportation in leaf veins share a common and continuous sieve system, the C supply and consumption in roots should be related to C transportation in leaves but not to *leaf dry mass per area*, a key leaf economics trait reflecting C production. This could explain our finding of no allometry of C production (i.e., *leaf dry mass per area*) with C supply (i.e., stele) and C consumption (i.e., *thickness of root tissues outside the stele*) in non-mangrove plants (Fig.3b).

Surprisingly, we found no correlation between C supply in the root stele and C transportation in leaf veins in mangrove plants, although roots and leaves share a continuous sieve system, while we did find a correlation between C supply in roots and C production leaves (Fig. 3a). These unexpected results can also be attributed to the presence of leaf *water storage tissues* that affect vein vessels and sieves in mangrove leaves differently. As mentioned previously, vein vessels in mangroves suffer from two opposing selection pressures with increasing salinity: positive pressure due to increased leaf *water storage tissues* and negative pressure due to increased physiological drought. In contrast to vessels, sieves in mangrove leaf veins do not seem to be markedly affected by leaf *water storage tissues* because of the storage of water rather than C (Fig. 4), and the sieves tended to be thin under increasing salinity stress to match the reduction in photosynthesis and C production^32,47^. Therefore, in contrast to non-mangrove plants (Fig. 3b), C production (represented by *leaf dry mass per area*) in mangrove leaves should be coupled with vein sieve-based C transportation, and consequently linked with root C supply (through sieves in the stele) and C consumption (through cells of *root tissues outside the stele thickness*) in an allometric manner (Fig. 3a). Root-leaf allometry paves a new way to understand the above- and belowground interactions regarding key components of C cycling within mangrove plants; that is, C production and transportation in leaves and C supply and consumption in roots.

In conclusion, our results revealed, for the first time, decoupled relationships between leaf hydraulics and economics and between root and leaf hydraulics, as well as allometry between *root tissues outside the stele* and the stele with a shift in *leaf dry mass per area*. In contrast, non-mangrove plants had leaf hydraulics coupled with root hydraulics while decoupled from leaf economics, and they also lacked leaf-root allometry. The contrasting leaf-leaf and leaf-root relationships could be due to the presence of leaf *water storage tissues* in mangroves that is absent from non-mangroves. Our study highlights the profound effects of leaf *water storage tissues* in shaping whole-plant strategies of resource (water and C) absorption, transportation, and consumption, potentially through its different impacts on vein vessels and sieves (Fig. 4). Give the widespread of leaf *water storage tissues* in mangroves, the novel leaf-root coordination in mangroves relative to non-mangroves potentially represents a new paradigm with which we look into the ecology, physiology and evolution of the important vegetation on Earth. In this sense, these findings are much insightful for understanding and prediction the dynamics and responses of mangrove ecosystems to environmental change. Last but not the least, our findings suggest that future studies on genetic improvement of mangrove plants should be oriented to identify key genetic factors or pathways governing interspecific variation of leaf *water storage tissues*.

## Materials and methods

### Sampling site and sample collection

This study was conducted in the Dongzhai Harbor National Natural Reserve (19°51′–20°01′N, 110°30′–110°37′E) in northeast Haikou City, Hainan province, China, where the mangrove ecosystem is located. It is the first and the largest mangrove reserve in China, with the most representative and pristine natural mangrove distribution. The area has a tropical monsoon climate. The mean annual temperature is 23.5 °C, with the highest mean monthly temperature, 28.4 °C, in July and the lowest, 17.1 °C, in January. The annual precipitation is 1,676 mm, and more than 80% is concentrated between May and October.

We collected the absorptive roots and leaves of 17 mangrove species typical at this reserve site in May 2015 (see Table S7 for details). Five mature trees were selected from each species. We selected 15–20 mature and intact canopy leaves and 3–5 intact root branches, including the first 3 terminal root orders for each individual. All root branches and a portion of leaves were immediately placed in FAA fixation solution (90 mL 70% alcohol, 5 mL 100% acetic acid, and 5 mL 37% methanol) for root and leaf anatomy measurements. Separate subsamples of the leaves were placed in NaOH solution to determine leaf vein diameter and density. The remaining leaf samples were used to measure for measurements of leaf morphology and chemicals.

### Trait measurements

We first determined *leaf thickness* for each species. The leaves were then scanned before the individual leaf area was calculated using IMAGE J software (NIH Image, Bethesda, MD, USA). The leaves were oven-dried at 60 °C for 48 h and were then weighed to determine *specific leaf area* (leaf area per unit leaf dry mass, SLA), *leaf dry mass per area* (LMA), and *leaf tissue density* (LTD). *Leaf mass-based nitrogen concentration* and *leaf C isotope composition* were measured using an elemental analyzer interfaced with isotope ratio mass spectrometry (EA1112 coupled with Delta-XP, Thermo Fisher Scientific, Bremen, Germany). The above leaf morphological and chemical traits are usually considered leaf economics traits (closely related to leaf photosynthesis). Leaf *total phenol content*, an indicator of leaf defense function, usually shows a trade-off with leaf photosynthesis^48^. Therefore, we considered *total phenol content* as a leaf economics trait, and which was measured by referring to the method in a classical study^49^.

Because mangrove leaves are usually thick and cuticle-rich for resistance to alkali decay, we first removed the epidermis and upper layers of the mesophyll to expose the leaf minor veins using a sharp knife or needle. The leaves were then dipped into a 5%–7% NaOH solution for hours to expose the leaf minor veins. The leaf veins were stained red and photographed using a camera (Eclipse Ni-U; Nikon). For each species, at least six fields of view per leaf were selected to calculate *leaf minor vein diameter* (LV_dia_, μm) and *density* (LV_den_, mm mm^-2^) using IMAGE J (NIH Image, Bethesda, MD, USA)^7^.

Leaf and root anatomical structures were determined using a common method for paraffin sectioning. Briefly, several leaves and the first-order roots were taken from the FAA solutions and were then processed in a suite of procedures, including dehydration, embeding in paraffin, cutting into sections (8 μm thickness), staineding and then photographing^5,26^. First-order root and leaf anatomical structures were determined using IMAGE J software (NIH Image, Bethesda, MD, USA). Specifically, we measured *root diameter, root stele diameter*, and *thickness of root tissues outside the stele* (including epidermis, exodermis and cortex), as these traits are closely related to water and nutrient absorption and transportation^18,27^. The *stele to root diameter ratio*, a key trait for root hydraulics, was also calculated^26^. Leaf *water storage tissues* have been widely observed in the hypodermis, sponge tissue and trichome layer in mangroves^14,15^. Here, we defined the size of leaf *water storage tissues* as the proportion of leaf cross-sectional area by the sum of the cross-sectional areas of the hypodermis, sponge tissue and trichome layer. In this study, leaf *water storage tissues* was an important trait, seemingly linked with both leaf economics and leaf hydraulics. To determine which trait spectra leaf *water storage tissues* belongs to, we first examined the correlations of leaf *water storage tissues* with leaf economics and leaf hydraulic traits separately. Then, the results of leaf *water storage tissues*-based pairwise correlations were further validated using a trait permutation analysis developed in a previous study^7^ (see Data analyses section for details).

To compare mangrove and non-mangrove species, we compiled a dataset from previous studies accessible for leaf hydraulic, economics, and root hydraulic traits. As most of the mangrove species in our study were woody, we selected only published studies on woody species^7,26^, and 78 species were included. We acknowledge that the leaf traits used in the comparing mangrove and non-mangrove plants were not the same. For example, we only had leaf vein traits and lacked leaf stomatal traits in mangrove plants. Nevertheless, incomplete mangrove leaf hydraulic traits could hardly markedly affect the relationships between leaf hydraulics and leaf economics and root hydraulics because of the universal coupling between leaf vein and stomatal traits^50,51,52,53,54^.

Additionally, only four leaf economics traits were only included in the mangrove plants: *specific leaf area*, leaf *water storage tissues, leaf tissue density* and *total phenol content*. Inclusion of *specific leaf area* should have little effect on the trait relationships for mangrove plants because *specific leaf area* is only a mathematical reciprocal of le*af dry mass per area*. Even after removing the three traits included only in the current mangrove study, that is, leaf *water storage tissues, leaf tissue density*, and *total phenol content*, the trait relationships within leaves and between leaves and roots were almost the same (see Fig. S8 and S9 in the Supporting Information). Therefore, it is feasible to compare mangrove and non-mangrove plants.

### Phylogenetic tree construction

Phylogenetic trees for the mangrove plants of this study and non-mangrove plants from previous studies^7,26^ (Fig. S10, S11) were constructed separately using the R package V. PhyloMaker^55,56^. Plant names in the phylogenetic tree were referenced against The Plant List (http://www.theplantlist.org/).

### Data analyses

We calculated the mean, minimum, maximum, standard error and coefficient of variation (CV) of each root and leaf trait. We calculated the phylogenetic signal by employing Blomberg’s *K* test, assuming a Brownian evolution model. A higher *K* value for a trait indicates more phylogenetic conservatism (i.e., more influence of the trait by a common ancestor than by environments)^57^. All the original data of non-mangrove plants and partial data of mangrove plants that did not confirm to a normal distribution, such as *root diameter, root stele diameter, thickness of root tissues outside the stele, leaf minor vein diameter and density, specific leaf area*, and *leaf mass-based nitrogen concentration*, were log-transformed. Pairwise trait relationships were assessed using Pearson’s correlation. We also explored trait relationships using phylogenetically independent contrasts (PICs), which exclude the influence of a common ancestor on trait relationships.

To explore the relationships between leaf hydraulics and economics in leaves, and the relationships between root hydraulics and leaf hydraulics and economics, we used a multivariate ordination method developed in a previous study^7^. All the data were standardized, and three separate PCAs were conducted using three different trait groups (i.e., leaf economics, leaf hydraulics, and root hydraulics) (Fig. 2a, b). Subsequently, we created a sampling distribution using the scores of 10,000 permutations of the first principal component axis (PC1) of each PCA. Finally, the relationships among leaf economics, leaf hydraulic and root hydraulic traits were assessed based on the permutation results^27^.

To test whether there was an allometric relationship between *root tissues outside the stele* and the stele in mangrove species, we compared the slopes of the regressions of *thickness of root tissues outside the stele* and stele radius with increasing root diameter using the standardized major axis method. Furthermore, if allometry between *root tissues outside the stele* and stele existed, we also explored whether within-root allometry could be applied to root-leaf allometry. Specifically, we first explored whether there were correlations between *leaf dry mass per area*, a key leaf economics trait, with *thickness of root tissues outside the stele* and stele radius. Where present, the slopes of the regressions of *thickness* of *root tissues outside the stele* and stele radius with *leaf dry mass per area*, were compared using the R package ‘smatr’. All data analyses were performed using R software (v. 3.30, R Core Team, 2016).

### Data availability

Data of mangrove species are available in Dryad Digital Repository, a publicly available database, or acquirable on request of the corresponding author. Data of root and leaf traits for non-mangrove species can be assessed in the supporting information of two previous studies^7,26^.

## Acknowledgements

We thank Mr. Lingqun Kong and Jianhai Chen for assistance in plant materials collection; Haiyan Zhang, Chao Guan, Xinyu Lu, Jinqi Tang, Qubing Ran, Song Huang and Mengke Wang for their help of trait measurements in the lab and phylogenetic analysis; and Editage (www.editage.cn) for proofreading manuscript. We are grateful to Dongzhai Harbor mangrove Wetland National Nature Reserve for their support. This study was funded by the National Natural Science Foundation of China (32171746, 31870522 and 31670550), and the Scientific Research Foundation of Henan Agricultural University (30500854), Research Funds for overseas returnee in Henan Province, China.

## Author contribution

D.K. and J.C. conceived the idea and collected the data. J.C., D.K. and W.R. conducted the statistical analyses. J.C., M.L. and D.K. wrote the draft of the manuscript. All authors contributed to the discussion of the results, manuscript revision and completion.

