## Supporting information for "Novel leaf-root coordination driven by leaf water storage tissues in mangroves"

**Table S1** Reports from principal components analysis on the10 leaf traits of 17 mangrove species, including the proportion of variation explained (top table) and loading scores of traits on each component (bottom table).

| Component | Eigenvalue | Proportion | Cumulative |
| --- | --- | --- | --- |
| 1 | 5.253 | 52.53% | 52.53% |
| 2 | 2.128 | 21.28% | 73.81% |
| 3 | 0.938 | 9.38% | 83.19% |
| 4 | 0.660 | 6.60% | 89.80% |
| 5 | 0.521 | 5.21% | 95.00% |
| 6 | 0.302 | 3.02% | 98.03% |
| 7 | 0.102 | 1.02% | 99.05% |
| 8 | 0.072 | 0.72% | 99.77% |
| 9 | 0.018 | 0.18% | 99.95% |
| 10 | 0.005 | 0.05% | 100.00% |

| Variable | Component 1 | Component 2 | Component 3 |
| --- | --- | --- | --- |
| *Leaf thickness* | 0.39 | -0.04 | 0.18 |
| *Leaf dry mass per area* | 0.41 | 0.20 | -0.06 |
| *Leaf mass-based nitrogen concentration* | -0.40 | -0.03 | 0.25 |
| *Leaf carbon isotope composition* | 0.05 | 0.44 | 0.76 |
| *Specific leaf area* | -0.40 | -0.14 | 0.19 |
| *Leaf tissue density* | 0.16 | 0.49 | -0.43 |
| *Total phenol content* | 0.31 | 0.01 | 0.27 |
| *Water storage tissues* | 0.32 | 0.07 | 0.13 |
| *Leaf minor vein diameter* | -0.24 | 0.51 | -0.14 |
| *Leaf minor vein density* | 0.27 | -0.50 | 0.01 |

**Table S2** Pearson correlation coeffificients (lower diagonal) and phylogenetically independent contrasts (upper diagonal) among 14 root and leaf functional traits for 17 mangrove species. Abbreviations of the traits are: LTh, *leaf thickness*; LMA, *leaf dry mass per area*; N_mass_, *leaf mass-based nitrogen concentration*; δ^13^C, *leaf carbon isotope composition*; SLA, *specific leaf area*; LTD, *leaf tissue density*; Tphol, *total phenol content*; WST, *water storage tissues*; LV_dia_, *leaf minor vein diameter*; LV_den_, *leaf minor vein density*; Diam, *root diameter*; Stele, *root stele diameter*; ToS, *thickness of root tissues outside the stele*; Stele : Diam, *stele to root diameter ratio*.

|  | LTh | LMA | N_mass_ | δ^13^C | SLA | LTD | Tphol | WST | LV_den_ | LVdia | Diam | ToS | Stele | Stele : Diam |
| --- | --- | --- | --- | --- | --- | --- | --- | --- | --- | --- | --- | --- | --- | --- |
| LTh |  | **0.85** | **-0.85** | 0.29 | **-0.82** | 0.09 | **0.80** | **0.55** | **-0.52** | **0.51** | **-0.64** | **-0.60** | **-0.69** | -0.12 |
| LMA | **0.85** |  | **-0.94** | 0.48 | **-0.95** | **0.58** | **0.83** | **0.68** | -0.31 | 0.35 | **-0.68** | **-0.65** | **-0.76** | -0.08 |
| N_mass_ | **-0.77** | **-0.86** |  | -0.45 | **0.88** | -0.42 | **-0.65** | **-0.72** | 0.40 | **-0.52** | **0.56** | **0.52** | **0.72** | 0.25 |
| δ^13^C | 0.17 | 0.24 | 0.05 |  | -0.10 | 0.24 | 0.23 | 0.34 | 0.27 | -0.34 | -0.27 | -0.27 | -0.21 | 0.02 |
| SLA | **-0.87** | **-0.94** | **0.90** | -0.24 |  | **-0.57** | **-0.68** | **-0.70** | 0.30 | -0.38 | **0.72** | **0.68** | **0.83** | 0.12 |
| LTD | 0.06 | **0.55** | -0.46 | 0.38 | -0.48 |  | 0.26 | 0.44 | 0.27 | -0.20 | -0.33 | -0.32 | -0.42 | 0.04 |
| Tphol | **0.52** | **0.61** | **-0.87** | **0.60** | **-0.51** | 0.24 |  | **0.59** | -0.39 | **0.48** | -0.37 | -0.35 | -0.42 | -0.14 |
| WST | **0.62** | **0.71** | **-0.60** | 0.19 | **-0.64** | 0.30 | 0.44 |  | -0.36 | 0.40 | **-0.69** | **-0.70** | **-0.52** | 0.21 |
| LV_den_ | **-0.63** | -0.41 | 0.41 | 0.32 | 0.45 | 0.11 | -0.39 | -0.16 |  | **-0.90** | 0.30 | 0.30 | 0.22 | -0.09 |
| LV_dia_ | **0.73** | 0.45 | **-0.52** | -0.30 | **-0.51** | -0.26 | 0.49 | 0.25 | **-0.89** |  | -0.38 | -0.38 | -0.35 | -0.02 |
| Diam | **-0.56** | -0.46 | **0.52** | 0.07 | **0.61** | -0.07 | -0.45 | **-0.74** | 0.43 | **-0.57** |  | **0.71** | **0.99** | -0.02 |
| ToS | **-0.64** | **-0.77** | **0.82** | -0.43 | **0.81** | -0.50 | **-0.70** | **-0.76** | 0.13 | -0.29 | **1.00** |  | **0.80** | **0.63** |
| Stele | **-0.51** | -0.39 | 0.46 | 0.14 | **0.55** | -0.04 | -0.39 | **-0.71** | 0.47 | **-0.58** | **0.85** | **0.64** |  | -0.12 |
| Stele : Diam | -0.39 | **-0.59** | **0.62** | **-0.82** | 0.43 | -0.43 | **-0.62** | -0.38 | -0.31 | 0.17 | -0.31 | -0.38 | 0.17 |  |

Significant correlations are indicated in bold.

Table S3 Reports from principal components analysis on the 8 leaf traits of 78 non-mangrove species, including the proportion of variation explained (top table) and loading scores of traits on each component (bottom table).

| Component | Eigenvalue | Proportion | Cumulative |
| --- | --- | --- | --- |
| 1 | 3.03 | 37.83% | 37.83% |
| 2 | 1.99 | 24.93% | 62.77% |
| 3 | 0.92 | 11.49% | 74.25% |
| 4 | 0.81 | 10.10% | 84.35% |
| 5 | 0.50 | 6.20% | 90.55% |
| 6 | 0.42 | 5.25% | 95.80% |
| 7 | 0.34 | 4.19% | 100.00% |
| 8 | 0.00 | 0.00% | 100.00% |

| Variable | Component 1 | Component 2 | Component 3 |
| --- | --- | --- | --- |
| *Leaf thickness* | -0.41 | 0.33 | 0.02 |
| *Leaf dry mass per area* | -0.30 | 0.40 | -0.36 |
| *Leaf mass-based nitrogen concentration* | 0.30 | -0.32 | 0.53 |
| *Leaf carbon isotope composition* | -0.15 | 0.43 | 0.59 |
| *Leaf minor vein density* | 0.38 | -0.05 | -0.31 |
| *Stomatal guard cell length* | -0.41 | -0.22 | 0.33 |
| *Stomatal density* | 0.44 | 0.44 | 0.02 |
| *Maximum stomatal conductance to water vapor* | 0.35 | 0.45 | 0.21 |

**Table S4** Pearson correlation coeffificients (lower diagonal) and phylogenetically independent contrasts (upper diagonal) among 12 root and leaf functional traits for 78 non-mangrove species. Abbreviations of the traits are: LTh, *leaf thickness*; LMA, *leaf dry mass per area*; N_mass_, *leaf mass-based nitrogen concentration*; δ^13^C, *leaf carbon isotope composition*; LV_den_, *leaf minor vein density*; SL, *stomatal guard cell length*; SD, *stomatal density*; g_wmax,_ *maximum stomatal conductance to water vapor*; Diam, *root diameter*; ToS, *thickness of root tissues outside the stele*; Stele, *root stele diameter*; Stele : Diam, *stele to root diameter ratio*.

|  | LTh | LMA | N_mass_ | δ^13^C | LV_den_ | SL | SD | g_wmax_ | Diam | ToS | Stele | Stele : Diam |
| --- | --- | --- | --- | --- | --- | --- | --- | --- | --- | --- | --- | --- |
| LTh |  | **0.61** | **-0.28** | **0.76** | **-0.82** | **0.79** | **-0.82** | **-0.80** | **0.80** | **0.76** | **0.51** | **-0.79** |
| LMA | **0.52** |  | -0.13 | **0.35** | **-0.39** | **0.56** | **-0.68** | **-0.68** | **0.45** | **0.35** | **0.40** | **-0.39** |
| N_mass_ | **-0.47** | **-0.49** |  | **-0.56** | **0.58** | **-0.26** | 0.17 | 0.13 | **-0.45** | **-0.56** | -0.10 | **0.55** |
| δ^13^C | **0.48** | **0.27** | -0.09 |  | **-0.95** | **0.72** | **-0.73** | **-0.70** | **0.98** | **1.00** | **0.66** | **-0.96** |
| LV_den_ | **-0.40** | -0.21 | **0.32** | -0.13 |  | **-0.73** | **0.76** | **0.74** | **-0.92** | **-0.95** | **-0.49** | **0.97** |
| SL | **0.29** | 0.11 | -0.16 | 0.03 | **-0.48** |  | **-0.88** | **-0.81** | **0.78** | **0.72** | **0.62** | **-0.71** |
| SD | **-0.26** | -0.07 | 0.10 | 0.13 | **0.36** | **-0.70** |  | **0.99** | **-0.82** | **-0.73** | **-0.59** | **0.77** |
| g_wmax_ | -0.17 | -0.03 | 0.03 | 0.19 | 0.19 | **-0.34** | **0.91** |  | **-0.80** | **-0.70** | **-0.55** | **0.76** |
| Diam | **0.25** | 0.14 | -0.06 | 0.07 | **-0.43** | **0.32** | -0.17 | -0.03 |  | **0.98** | **0.73** | **-0.94** |
| ToS | **0.23** | 0.11 | -0.03 | 0.08 | **-0.43** | **0.32** | -0.16 | -0.03 | **0.99** |  | **0.66** | **-0.96** |
| Stele | **0.22** | 0.16 | -0.03 | 0.03 | **-0.33** | **0.26** | -0.13 | -0.02 | **0.95** | **0.92** |  | **-0.46** |
| Stele : Diam | **-0.24** | -0.05 | 0.11 | -0.19 | **0.49** | **-0.31** | 0.18 | 0.06 | **-0.65** | **-0.70** | **-0.39** |  |

Significant correlations are indicated in bold.

**Table S5** Reports from principal components analysis on the 4 root traits and 10 leaf traits of 17 mangrove species, including the proportion of variation explained (top table) and loading scores of traits on each component (bottom table).

| Component | Eigenvalue | Proportion | Cumulative |
| --- | --- | --- | --- |
| 1 | 7.102 | 50.73% | 50.73% |
| 2 | 2.221 | 15.87% | 66.60% |
| 3 | 1.632 | 11.66% | 78.26% |
| 4 | 0.928 | 6.63% | 84.89% |
| 5 | 0.783 | 5.59% | 90.48% |
| 6 | 0.530 | 3.79% | 94.26% |
| 7 | 0.349 | 2.49% | 96.76% |
| 8 | 0.285 | 2.03% | 98.79% |
| 9 | 0.078 | 0.56% | 99.35% |
| 10 | 0.054 | 0.39% | 99.74% |
| 11 | 0.020 | 0.14% | 99.88% |
| 12 | 0.013 | 0.09% | 99.97% |
| 13 | 0.004 | 0.03% | 100.00% |
| 14 | 0.000 | 0.00% | 100.00% |

| Variable | Component 1 | Component 2 | Component 3 |
| --- | --- | --- | --- |
| *Leaf thickness* | -0.33 | -0.11 | -0.09 |
| *Leaf dry mass per area* | -0.35 | 0.11 | -0.15 |
| *Leaf mass-based nitrogen concentration* | 0.33 | 0.07 | 0.28 |
| *Leaf carbon isotope composition* | -0.07 | 0.42 | 0.05 |
| *Specific leaf area* | 0.35 | -0.06 | 0.16 |
| *Leaf tissue density* | -0.15 | 0.43 | -0.15 |
| *Total phenol content* | -0.24 | -0.09 | -0.20 |
| *Water storage tissues* | -0.29 | 0.03 | 0.20 |
| *Leaf minor vein diameter* | 0.18 | 0.52 | -0.15 |
| *Leaf minor vein density* | -0.21 | -0.52 | 0.07 |
| *Root diameter* | 0.32 | -0.13 | -0.32 |
| *Thickness of root tissues outside the stele* | 0.31 | -0.13 | -0.37 |
| *Root stele diameter* | 0.32 | -0.12 | 0.06 |
| *Stele to root diameter ratio* | 0.00 | 0.07 | 0.70 |

Table S6 Reports from principal components analysis on the 12 root and leaf traits of 78 non-mangrove species, including the proportion of variation explained (top table) and loading scores of traits on each component (bottom table).

| Component | Eigenvalue | Proportion | Cumulative |
| --- | --- | --- | --- |
| 1 | 4.35 | 36.25% | 36.25% |
| 2 | 2.25 | 18.72% | 54.96% |
| 3 | 1.96 | 16.36% | 71.32% |
| 4 | 0.93 | 7.77% | 79.09% |
| 5 | 0.80 | 6.67% | 85.77% |
| 6 | 0.58 | 4.81% | 90.57% |
| 7 | 0.42 | 3.48% | 94.05% |
| 8 | 0.39 | 3.27% | 97.32% |
| 9 | 0.31 | 2.58% | 99.90% |
| 10 | 0.01 | 0.08% | 99.97% |
| 11 | 0.00 | 0.02% | 100.00% |
| 12 | 0.00 | 0.00% | 100.00% |

| Variable | Component 1 | Component 2 | Component 3 |
| --- | --- | --- | --- |
| *Leaf thickness* | 0.27 | -0.20 | 0.41 |
| *Leaf dry mass per area* | 0.18 | -0.14 | 0.46 |
| *Leaf mass-based nitrogen concentration* | -0.15 | 0.22 | -0.41 |
| *Leaf carbon isotope composition* | 0.12 | 0.03 | 0.44 |
| *Leaf minor vein density* | -0.33 | 0.12 | -0.05 |
| *Stomatal guard cell length* | 0.29 | -0.29 | -0.16 |
| *Stomatal density* | -0.24 | 0.50 | 0.30 |
| *Maximum stomatal conductance to water vapor* | -0.14 | 0.49 | 0.31 |
| *Root diameter* | 0.41 | 0.30 | -0.11 |
| *Thickness of root tissues outside the stele* | 0.41 | 0.31 | -0.13 |
| *Root stele diameter* | 0.37 | 0.31 | -0.10 |
| *Stele to root diameter ratio* | -0.34 | -0.16 | 0.07 |

**Table S7** Summary information for the 17 mangrove plant species in study

| Code | Species | Genus | Family | Life form | Plant height (m) |
| --- | --- | --- | --- | --- | --- |
| 1 | *Acrostichum aureum* | *Acrostichum* | *Acrostichaceae* | Herbaceous | 0.5-1m |
| 2 | *Bruguiera sexangula* | *Bruguiera* | *Rhizophoraceae* | Small tree | 5-6m |
| 3 | *Ceriops tagal* | *Ceriops* | *Rhizophoraceae* | Small tree | 2-3m |
| 4 | *Rhizophora mangle* | *Rhizophora* | *Rhizophoraceae* | Small tree | 2-3m |
| 5 | *Laguncularia racemosa* | *Laguncularia* | *Combretaceae* | Tree | 6-7m |
| 6 | *Bruguiera gymnorrhiza* | *Bruguiera* | *Rhizophoraceae* | Small tree | 3-4m |
| 7 | *Rhizophora mucronata* | *Rhizophora* | *Rhizophoraceae* | Small tree | 4-5m |
| 8 | *Rhizophora stylosa* | *Rhizophora* | *Rhizophoraceae* | Small tree | 3-4m |
| 9 | *Avicennia marina* | *Avicennia* | *Verbenaceae* | Small tree | 2-3m |
| 10 | *Kandelia candel* | *Kandelia* | *Rhizophoraceae* | Small tree | 3-4m |
| 11 | *Excoecaria agallocha* | *Excoecaria* | *Euphorbiaceae* | Small tree | 3-4m |
| 12 | *Sonneratia apetala* | *Sonneratia* | *Sonneratiaceae* | Tree | 7-8m |
| 13 | *Aegiceras corniculatum* | *Aegiceras* | *Myrsinaceae* | Shrub | 1.5-2m |
| 14 | *Ceriops tagal* | *Ceriops* | *Rhizophoraceae* | Shrub | 1-1.5m |
| 15 | *Thespesia populnea* | *Thespesia* | *Malvaceae* | Tree | 5-6m |
| 16 | *Pongamia pinnata* | *Pongamia* | *Leguminosae* | Tree | 5-6m |
| 17 | *Heritiera littoralis* | *Heritiera* | *Sterculiaceae* | Tree | 10-12m |

**Figure S1** Observed correlations (vertical lines) relative to the distribution of 17 mangrove species (a) when scores were randomly simulated between PC1 scores of *water storage tissues*(WST) and PC1 scores of leaf economics traits (*leaf thickness*, *specific leaf area*, *leaf dry mass per area*, *leaf tissue density*, *total phenol content*, *leaf mass-based nitrogen concentration*, *leaf carbon isotope composition*). The correlation is statistically significant with *p* = 0.003, demonstrating leaf *water storage tissues*(WST) coupled with leaf economics (LES); (b) when scores were randomly simulated between PC1 scores of WST and PC1 scores of leaf hydraulic traits (including *leaf minor vein diameter*, *leaf minor vein density*). The correlation is statistically significant with *p* = 0.338, demonstrating leaf *water storage tissues* (WST) are orthogonal to leaf hydraulics (LHS).
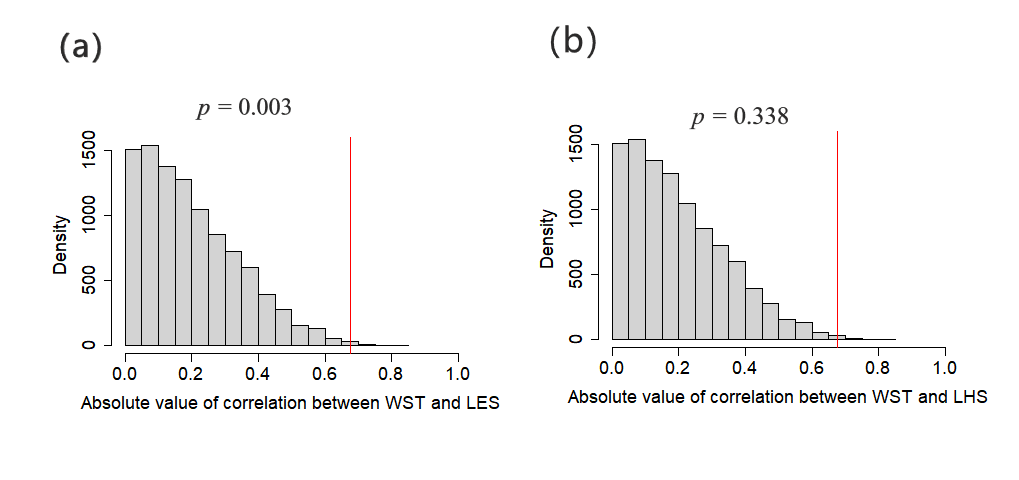

**Figure S2** Observed correlations (vertical lines) relative to the distribution of 17 mangrove species when scores of the principal component analysis (PCA) were randomly simulated (a) between PC1 (i.e., the first principal component of the PCA) scores of leaf hydraulic traits (including *leaf minor vein diameter*, *leaf minor vein density*; colored in green in Fig.1a) and PC1 scores of leaf economics traits (including *leaf thickness*, *specific leaf area*, *leaf dry mass per area*, *leaf tissue density*, *total phenol content*, *leaf mass-based nitrogen concentration*, *leaf carbon isotope composition*, *water storage tissues*; colored in blue in Fig.1a); (b) between PC1 scores of root hydraulic traits and PC1 scores of leaf hydraulic traits; (c) between PC1 scores of root hydraulic traits (*root diameter*, *thickness of root tissues outside the stele*, *root stele diameter*, *stele to root diameter ratio*; colored in red in Fig.1a) and PC1 scores of leaf economics traits. The correlations of (a) and (b) are non-significant with *p* values of 0.100 and 0.182, demonstrating that leaf hydraulics are orthogonal to leaf economics, and root hydraulics is orthogonal to leaf hydraulics. But the correlation of (c) is significant with *p* < 0.001, demonstrating that root hydraulics is coupled with leaf economics. Abbreviations: LHS, leaf hydraulics; LES, leaf economics; RHS, root hydraulics.

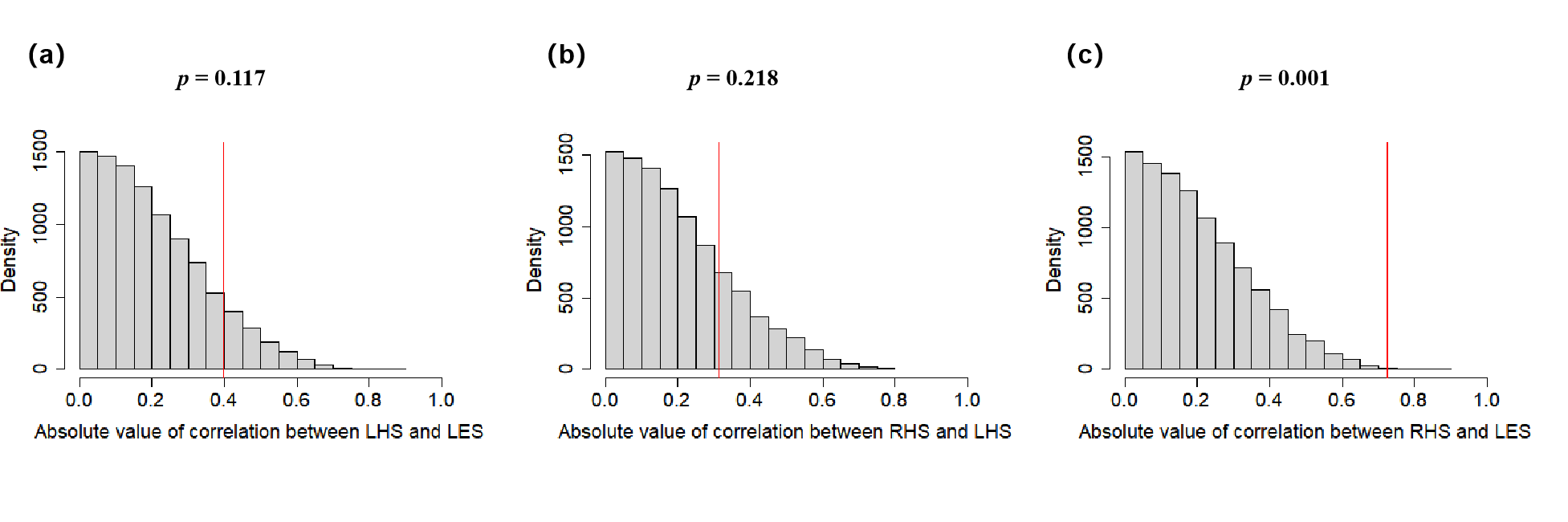

**Figure S3** Observed correlations (vertical lines) relative to the distribution of 78 non-mangrove species when scores of the principal component analysis (PCA) were randomly simulated (a) between PC1 (i.e., the first principal component of the PCA) scores of leaf hydraulic traits (including *leaf minor vein density*, *stomatal guard cell length*, *stomatal density*, *maximum stomatal conductance to water vapor*; colored in green in Fig.1b) and PC1 scores of leaf economics traits (including *leaf thickness*, *leaf dry mass per area*, *leaf mass-based nitrogen concentration*, *leaf carbon isotope composition*; colored in blue in Fig.1b); (b) between PC1 scores of root hydraulic traits and PC1 scores of leaf hydraulic traits; (c) between PC1 scores of root hydraulic traits (*root diameter*, *thickness of root tissues outside the stele*, *root stele diameter*, *stele to root diameter ratio*; colored in red in Fig.1b) and PC1 scores of leaf economics traits. The correlations of (a) and (c) are non-significant with *p* values of 0.062 and 0.073, demonstrating that leaf hydraulics is orthogonal to leaf economics, and root hydraulics is orthogonal to leaf economics. But the correlation of (b) is significant with *p* values of 0.012, demonstrating that root hydraulics is coupled with leaf hydraulics. Abbreviations: LHS, leaf hydraulics; LES, leaf economics; RHS, root hydraulics.

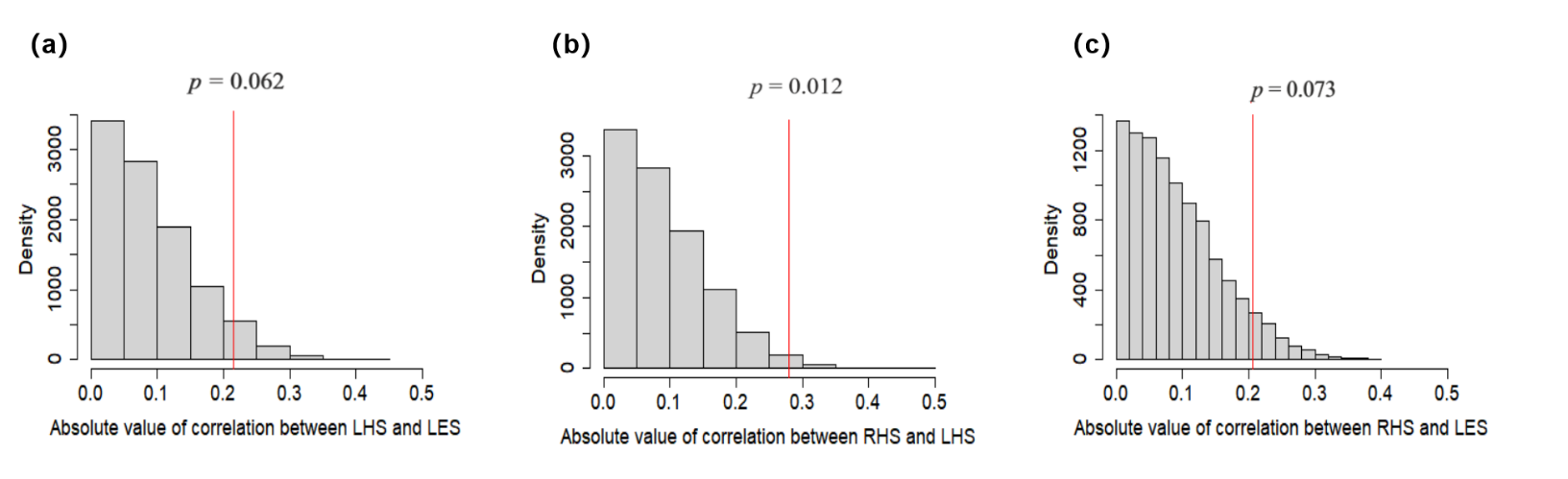

**Figure S4** Principal component analysis (PCA) for the 14 leaf and root traits of 17 mangrove species using phylogenetic independent contrasts (PICs). The leaf hydraulics traits are in green, leaf economics traits are in blue and root hydraulics traits are in red. Abbreviations of the traits are: LTh, *leaf thickness*; LMA, *leaf dry mass per area*; N_mass_, *leaf mass-based nitrogen concentration*; δ^13^C, *leaf carbon isotope composition*; SLA, *specific leaf area*; LTD, *leaf tissue density*; Tphol, *total phenol content*; WST, *water storage tissues*; LV_dia_, *leaf minor vein diameter*; LV_den_, *leaf minor vein density*; Diam, *root diameter*; ToS, *thickness of root tissues outside the stele*; Stele, *root stele diameter*; Stele : Diam, *stele to root diameter ratio*.

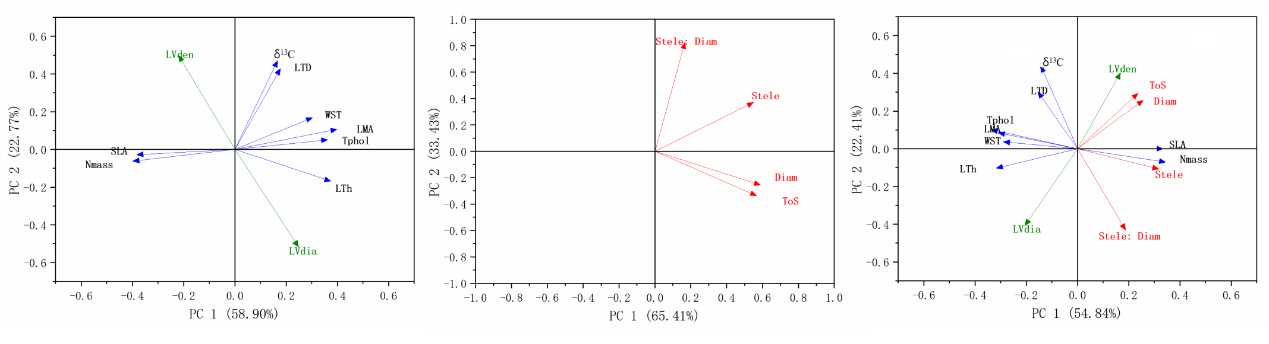

**Figure S5** Observed correlations (vertical lines) relative to the distribution of 17 mangrove species using phylogenetic independent contrasts (PICs) when scores were randomly simulated (a) between PC1 scores of leaf hydraulic traits (including *leaf minor vein diameter*, *leaf minor vein density*; colored in green in Fig.1a) and PC1 scores of leaf economics traits (including *leaf thickness*, *specific leaf area*, *leaf dry mass per area*, *leaf tissue density*, *total phenol content*, *leaf mass-based nitrogen concentration*, *leaf carbon isotope composition*, *water storage tissues*; colored in blue in Fig.1a); (b) between PC1 scores of root hydraulic traits and PC1 scores of leaf hydraulic traits; (c) between PC1 scores of root hydraulic traits (*root diameter*, *thickness of root tissues outside the stele*, *root stele diameter*, *stele to root diameter ratio*; colored in red in Fig.1a) and PC1 scores of leaf economics. The correlations of (a) and (b) are non-significant with *p* values of 0.111 and 0.105, demonstrating that leaf hydraulics (LHS) is orthogonal to leaf economics (LES), and root hydraulics (RHS) is orthogonal to leaf hydraulics (LHS). But the correlation of (c) are significant with *p* = 0.002, demonstrating that root hydraulic traits (RHS) is coupled with leaf economics (LES).

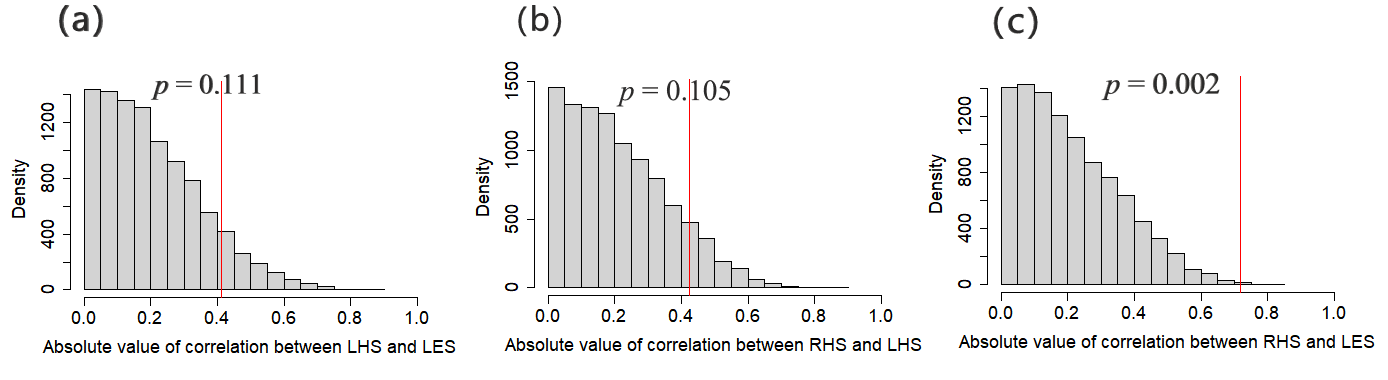

**Figure S6** Allometric relationships between root stele radius and *thickness of root tissues outside the stele* with increasing root diameter in mangrove (a) and non-mangrove species (b). Abbreviations: ToS, *thickness of root tissues outside the stele*; Diam, *root diameter*; LMA, *leaf dry mass per area*.

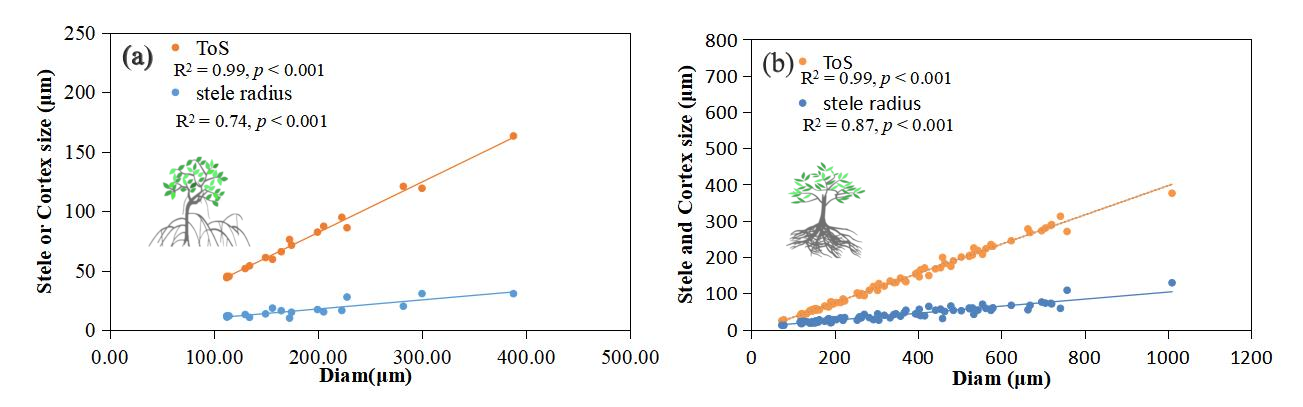

**Figure S7** Allometry between roots and leaves across 17 mangrove species. Abbreviations: LTh, *leaf thickness*; SLA, *specific leaf area*; N_mass_, *leaf mass-based nitrogen concentration*; WST, *water storage tissues*; ToS, *thickness of root tissues outside the stele*.

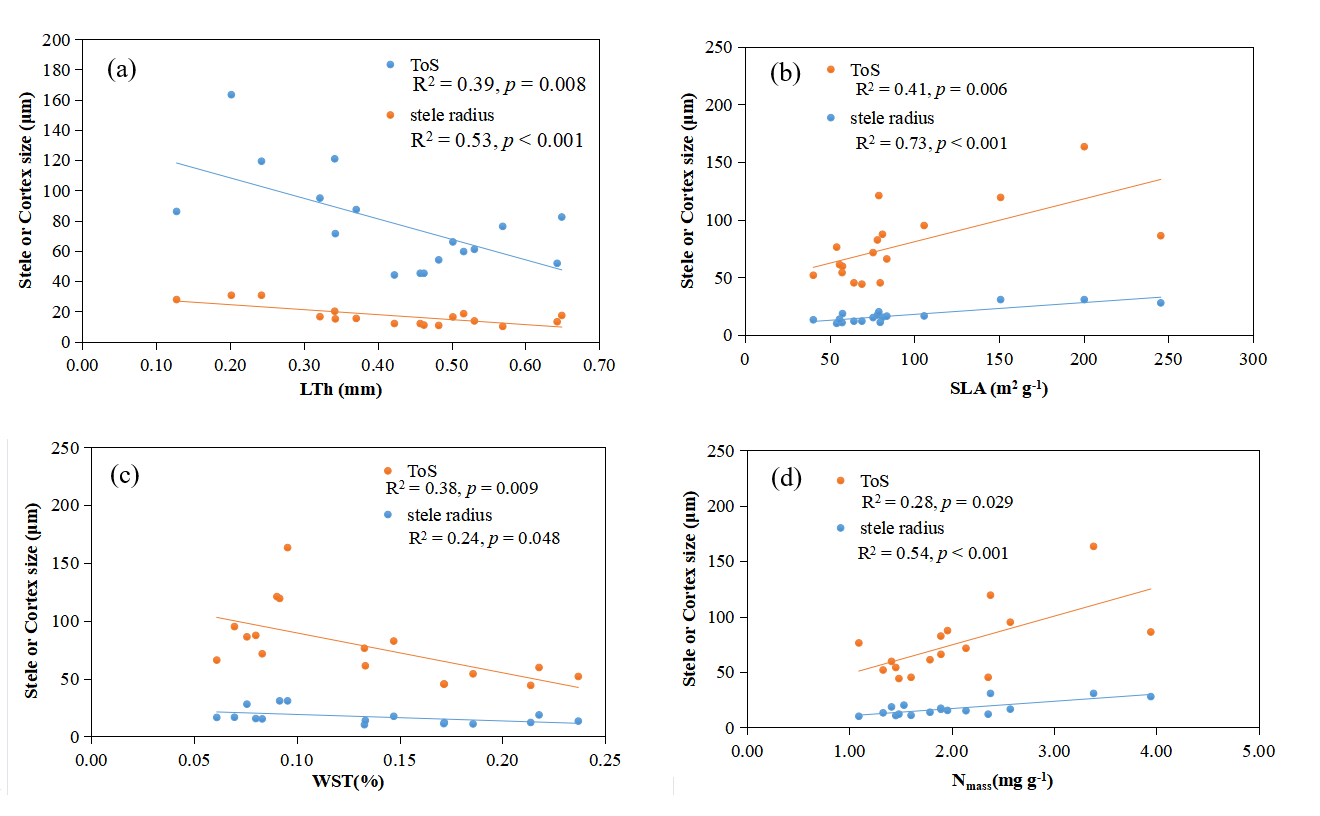

**Figure S8** Traits decoupling of leaf and root traits among mangrove plants after removing leaf water storage tissues, leaf tissue density and total phenol content. Traits abbreviation: LTh, *leaf thickness*; LMA, *leaf dry mass per area*; N_mass_, *leaf mass-based nitrogen concentration*; δ^13^C, *leaf carbonisotope composition*; SLA, *specific leaf area*; LV_dia_, *leaf minor vein diameter*; LV_den_, *leaf minor vein density*; Diam, *root diameter*; ToS, *thickness of root tissues outside the stele*; Stele, *root stele diameter*; Stele : Diam, *stele to root diameter ratio*.

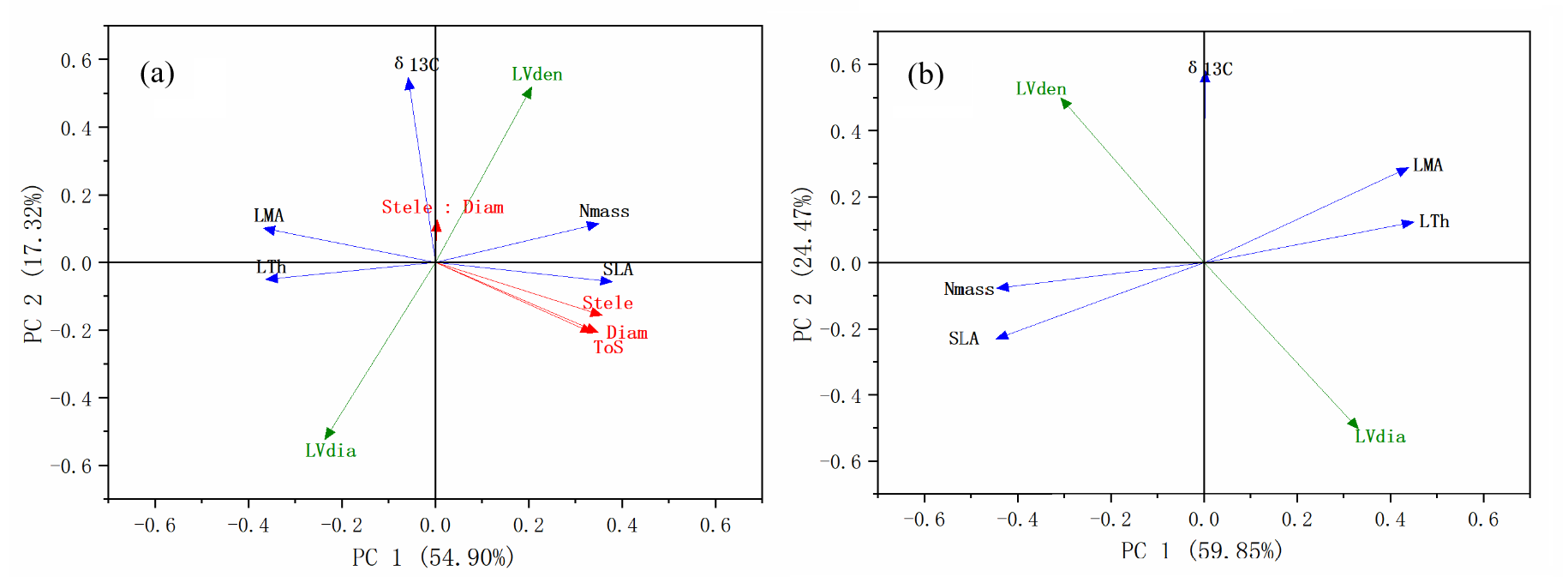

**Figure S9** Observed correlations (vertical lines) relative to the distribution of 17 mangrove species when scores of the principal component analysis (PCA) were randomly simulated as in Figure S5 after removing *leaf water storage tissues*, *leaf tissue density* and *total phenol content*.

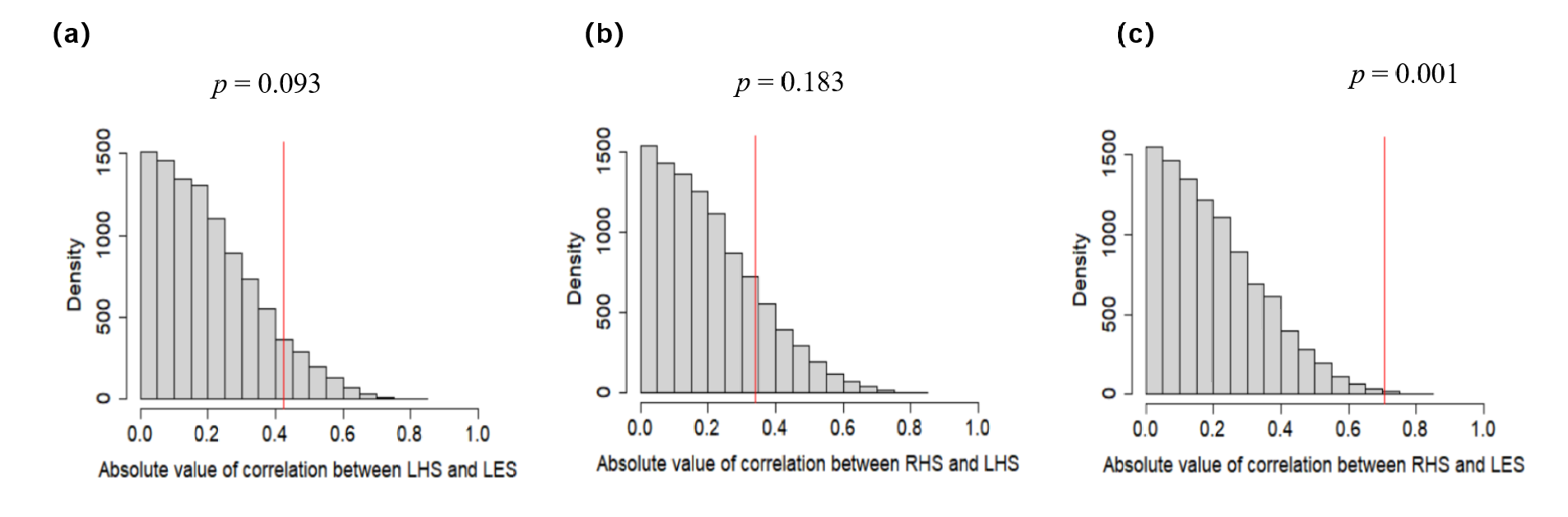

**Figure S10** Phylogenetic tree of 17 mangrove species. The size of the symbols is proportional to the difference of the traits from their average values across all species. Abbreviations of the traits are: LTh, *leaf thickness*; LMA, *leaf dry mass per area*; N_mass_, *leaf mass-based nitrogen concentration*; δ^13^C, *leaf carbon isotope composition*; SLA, *specific leaf area*; LTD, *leaf tissue density*; Tphol, *total phenol content*; WST, *water storage tissues*; LV_dia_, *leaf minor vein diameter*; LV_den_, *leaf minor vein density*; Diam, *root diameter*; ToS, *thickness of root tissues outside the stele*; Stele, *root stele diameter*; Stele : Diam, *stele to root diameter ratio*.

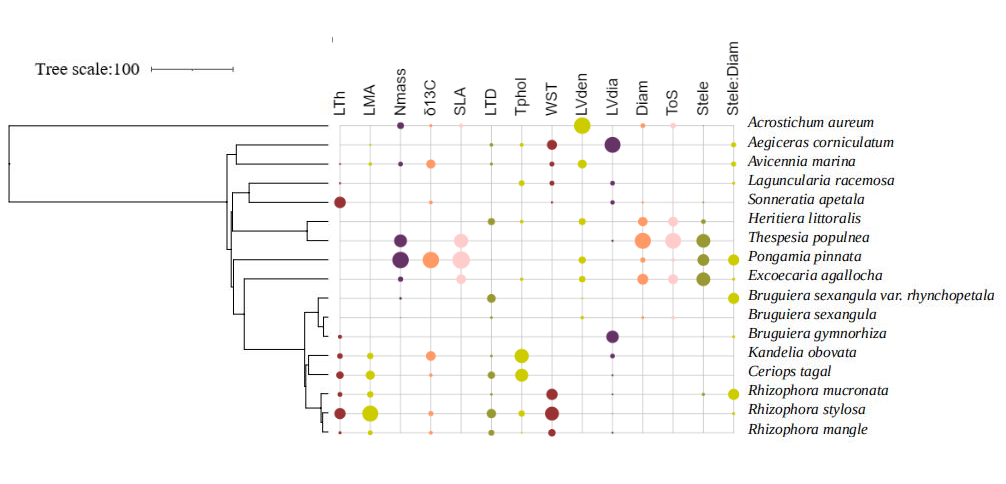

**Figure S11** Phylogenetic tree of 78 non-mangrove species.

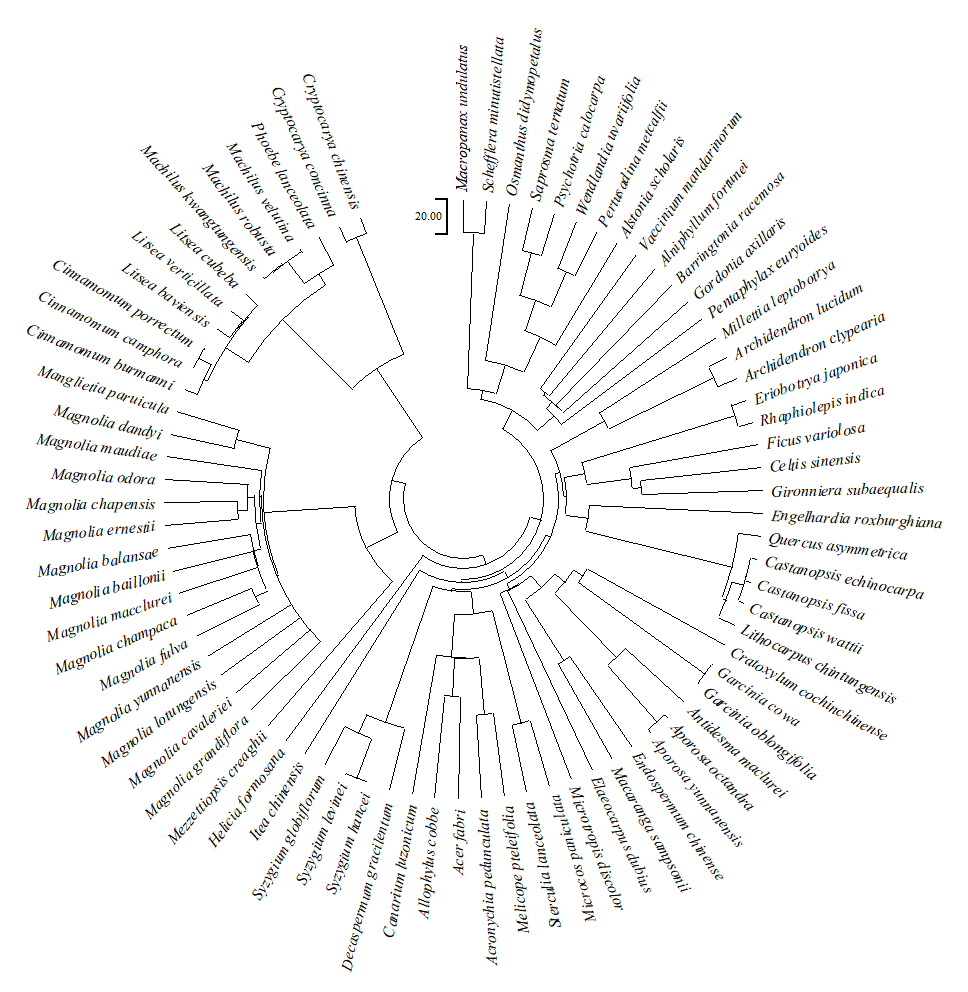
